# Single-cell RNA-Seq reveals the link between *CD45* isoforms and tumor-infiltrating T cells heterogeneity in liver cancer

**DOI:** 10.1101/2020.03.22.002824

**Authors:** Biaofeng Zhou, Shang Liu, Liang Wu, Yan Sun, Jie Chen, Shiping Liu

## Abstract

*CD45* isoforms play a major role in characterizing T cell function, phenotype, and development. However, there is lacking comprehensive interrogation about the relationship between *CD45* isoforms and T lymphocytes from cancer patients at the single-cell level yet. Here, we investigated the CD45 isoforms component of published 5,063 T cells of hepatocellular carcinoma (HCC), which has been assigned functional states. We found that the distribution of *CD45* isoforms in T lymphocytes cells depended on tissue resource, cell type, and functional state. Further, we demonstrated that CD45RO and CD45RA dominate in characterizing the phenotype and function of T cell though multiple *CD45* isoforms coexist in T cells, through a novel alternative splicing pattern analysis. We identified a novel development trajectory of tumor-infiltrating T cells from Tcm to Temra (effector memory T cells re-expresses CD45RA) after detecting two subpopulations in state of transition, Tcm (central memory T) and Tem (effector memory T). Temra, capable of high cytotoxic characteristics, was discovered to be associated with the stage of HCC and may be a target of immunotherapy. Our study presents a comprehension of the connection between *CD45* isoforms and the function, states, sources of T lymphocytes cells in HCC patients at the single-cell level, providing novel insight for the effect of *CD45* isoforms on T cell heterogeneity.

## 1. Introduction

*CD45*, a canonical marker for immune cell, presents various of isoforms arising from alternative splicing of the exons 4, 5 and 6 (corresponding to A, B, C) on all differentiated hematopoietic cells which showed cell-type and differentiation-stage specific expression(Hermiston, Xu, & Weiss, 2003; Holmes, 2006)(Hermiston et al. 2003; Holmes 2006) (Fig. 1A). For example, CD45RA and CD45RO are widely used to mark naïve T and memory T cells respectively. In addition, T cells can be divided into four types according to the expression of CD45RA and *CCR7*: Tnaive (naïve T cell, CD45RA+*CCR7*+), Tcm (central memory T, CD45RA-*CCR7*+), Tem (effector memory T, CD45RA-*CCR7*−) and Temra (effector memory T cells re-expresses CD45RA, CD45RA+*CCR7*−)(Sallusto, Lenig, Forster, Lipp, & Lanzavecchia, 1999). A relatively high number of Temra in peripheral blood was associated with good prognosis in NSCLC (non-small lung cancer) patients treated with nivolumab(Kunert et al., 2019). These indicate the vital importance of *CD45* isoforms in characterizing different types of T lymphocytes and prognosis in the cancer patients. However, almost all the previous studies about exploring the characterization and distribution of CD45 isoforms in T cells was done at bulk level with potential cell heterogeneity, which may affect the accuracy of results. For example, bimodality in splicing resulting in isoforms heterogeneity was revealed in immune cell with scRNA-seq while bulk RNA-seq fails(Shalek et al., 2013). Therefore, analyzing the distribution of *CD45* isoforms among T cells at single-cell level would deepen our understanding about the relationship between cell state and *CD45* isoforms.

**Figure 1.**
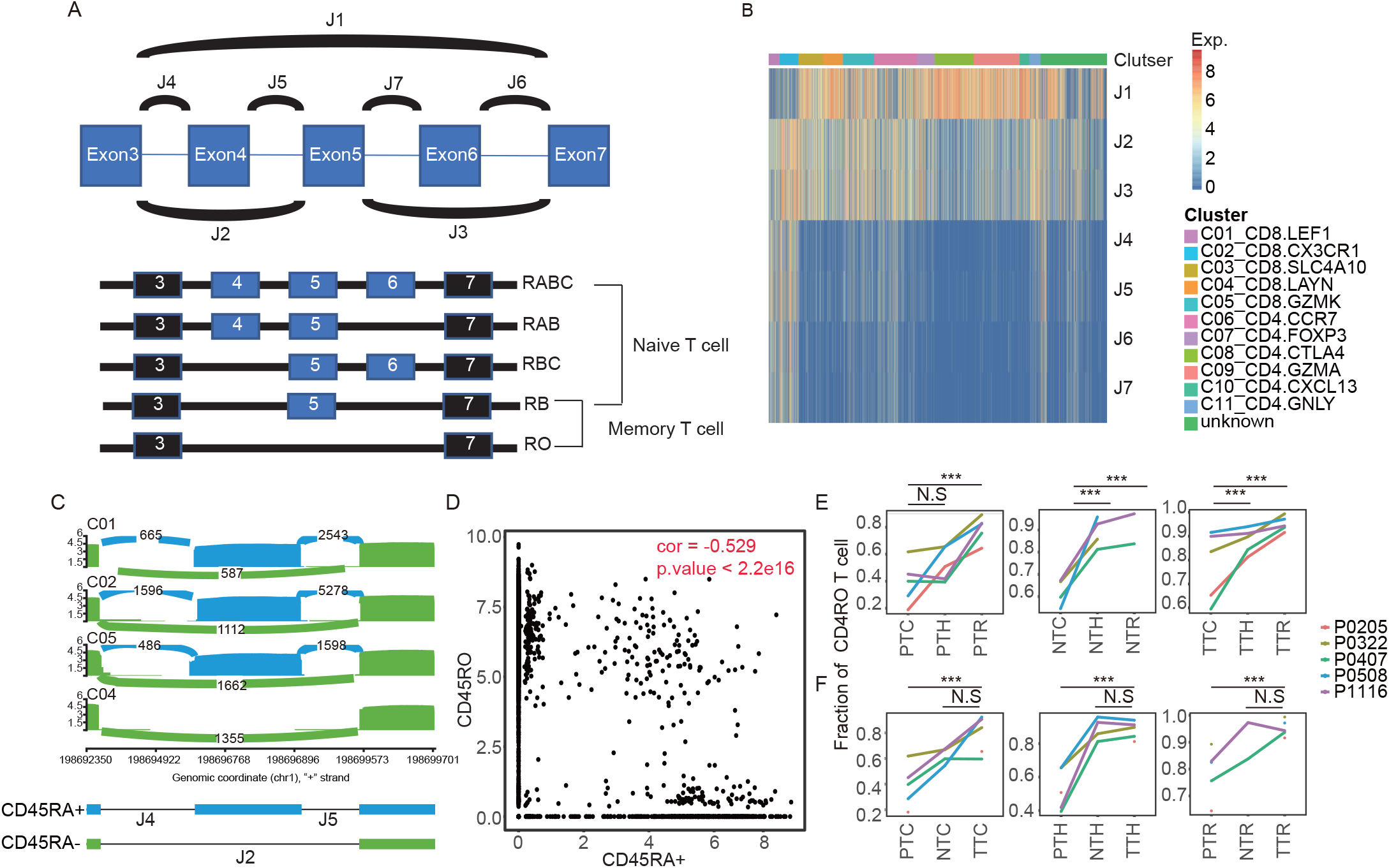
The distribution of CD45 isoforms of T cell in TME. (A) The carton shows the junction composition of five CD45 isoforms universally expressing in T cells. (B) The expression heat map of junctions related with five CD45 isoforms (see Fig. 5A). Information of pattern with different clusters are colored for each cell. (C) Sashimi plots illustrating the read distribution of PTPRC in CD8+ T cells from P0508 patient. Exhausted T cells (C04_CD8.LAYN) merely express CD45RA. (D) The expression of CD45RA and CD45RO across all T cells. (E) The percentages of cells expressing CD45RO (the normalized count of CD45RO > 1) in adjacent normal tissues, tumor tissues and blood. *p < 0.05, **p < 0.01, ***p < 0.001, N.S., not significant, Student’s t test. (F) The percentages of cells expressing CD45RO in cytotoxicity T cells, helper T cells and Tregs. *p < 0.05, **p < 0.01, ***p < 0.001, N.S., not significant, Student’s t test. Note: PTC, NTC, and TTC: CD8+ cytotoxic T cells from peripheral blood, adjacent normal, and tumor tissues; PTH, NTH, and TTH: T helpers cell from peripheral blood, adjacent normal, and tumor tissues; PTR, NTR, and TTR: Treg from peripheral blood, adjacent normal, and tumor tissues.

scRNA-seq has revolutionized our understanding about the heterogeneity of tumor infiltrating lymphocytes (TIL), since systematic interrogation of tumor-infiltrating lymphocytes were fulfilled in liver(Zheng et al., 2017), lung(Guo et al., 2018), colon(Zhang et al., 2018), and breast cancers(Chung et al., 2017). The comprehension of heterogeneity makes a great contribution to the identification and characterization of various types of TIL functional clusters such as exhausted T cell, naïve T cell, effector T, regulated T and so on. Besides, full length scRNA-seq data is also capable of isoform expression quantification. Those make it possible to research *CD45* isoforms distribution in high resolution.

Here, we systemically investigated the *CD45* isoforms component of published 5,063 T cells of hepatocellular carcinoma (HCC) from 6 patients. These T cells have been assigned cell types, functional states, and tissue resources. We discovered the distribution characterization of *CD45* isoforms as well as the dominant isoforms, CD45RO and CD45RA in T cell phenotype identification at single-cell level. We identified a novel development trajectory of tumor-infiltrating T cell from Tcm to Temra. Meanwhile, we found Temra with high cytotoxic characteristic, was related to HCC staging and may be a target of immunotherapy. All the results promote our understanding about the influence of *CD45* isoform on T cell heterogeneity.

## 2. Materials and methods

### 2.1 Data Sets

RNA-seq data of Human T cells in Fastq format was downloaded from EGD database (EGAS00001002072) after the access of the dataset has been approved. Matrix of gene expression was downloaded from GEO database with accession numbers GSE98638(https://www.ncbi.nlm.nih.gov/geo/query/acc.cgi?acc=GSE98638). It comprised 5063 T cells of 12 clusters from peripheral blood, tumor and adjacent normal liver tissue. The detailed clinical information of patient and cluster information was given in Table 1 and Table 2. The data was the result of single cell RNA sequencing by Illumina HiSeq2500 or Illumina Hiseq 4000. Necessary metadata was provided by article of Zheng’s(Zheng et al., 2017).

**Table 1.** The information of patients.

| Patient | Age | Sex | Stage | Tissue |
| --- | --- | --- | --- | --- |
| P1202 | 26 | Female | I | P/N/T |
| P0407 | 63 | Male | I | P/N/T |
| P0205 | 53 | Male | I | P/N/T |
| P0508 | 28 | Male | II | P/N/T/J |
| P0322 | 64 | Male | IVB | P/N/T |
| P1116 | 52 | Female | IVB | P/N/T |

Note: P: Peripheral blood, N: Normal Tissue, T: Tumor Tissue, J: adjacent normal tissues

**Table2.** Annotation about cell clusters

| Cluster | Cell number | Function annotation | Type |
| --- | --- | --- | --- |
| C01_CD8.LEF1 | 161 | Naïve T cell | CD8+ T cell |
| C02_CD8.CX3CR1 | 288 | Effector T cell | CD8+ T cell |
| C03_CD8.SLC4A10 | 363 | MAIT | CD8+ T cell |
| C04_CD8.LAYN | 300 | Exhausted T cell | CD8+ T cell |
| C05_CD8.GZMK | 467 | T cell in mediate state | CD8+ T cell |
| C06_CD4.CCR7 | 646 | Naïve T cell | CD4+ T cell |
| C07_CD4.FOXP3 | 261 | Peripheral Treg | CD4+ T cell |
| C08_CD4.CTLA4 | 582 | Tumor Treg | CD4+ T cell |
| C09_CD4.GZMA | 689 | T cell in mediate state | CD4+ T cell |
| C10_CD4.CXCL13 | 146 | Exhausted T cell | CD4+ T cell |
| C11_CD4.GNLY | 167 | Effector T cell | CD4+ T cell |
| Unknown | 993 | NA | NA |

### 2.2 Preprocessing of RNA-Seq data

Firstly, cleaned reads of each cell were aligned onto the UCSC hg38 human genome by STAR[10] with default parameters. Secondly, we obtained the read count of seven junctions of CD45 that involved in alternative splicing of *CD45*(Fig 1A) from the SJ.out.tab files of each cell. Thirdly, the read count of seven junctions was normalized by the uniquely mapped reads number of each cell. Finally, we obtained a junction-cell matrix with seven junctions as row and 5,036 cells as column.

### 2.3 Cell filtering

we used K-means to cluster cells with the normalized count and then filtered the cell clusters whose mean and standard deviation of normalized junction read count was less than 1 and 0.2 respectively. These filter cells could not reflect the real junction composition of *CD45* because of the low sequencing depth and high dropout rate in single-cell RNA-seq. At last, 79 cells which satisfied the criteria were filtered. Sashimi plots of gene were generated using the software package MISO(Katz, Wang, Airoldi, & Burge, 2010)

### 2.4 Junction expression pattern analysis of CD45

We first filtered cells in unknown clusters and then defined highly expressed junctions as log2 (normalized read count + 1) > 2.5 for those seven junctions of *CD45*. Finally, a total of 3,524 cells were assigned into nine major patterns after filtering. The isoforms component of each pattern is visible in Table 3.

**Table 3.** CD45 Isoform component of each pattern

| Pattern | J1 | J2 | J3 | J4 | J5 | J6 | J7 | component |
| --- | --- | --- | --- | --- | --- | --- | --- | --- |
| Pattern1 | - | - | - | + | + | + | + | RABC |
| Pattern2 | - | + | + | + | + | + | + | unclarity |
| Pattern3 | - | + | - | + | + | + | + | RABC+RBC |
| Pattern4 | - | + | + | - | - | - | - | RB |
| Pattern5 | + | + | + | - | - | - | - | RO+RB |
| Pattern6 | + | - | - | - | - | - | - | RO |
| Pattern7 | - | + | + | + | + | - | - | RB+RAB |
| Pattern8 | - | + | + | - | - | + | + | RBC+RB |
| Pattern9 | + | + | + | - | - | + | + | RO+RBC+RB |

Unclarity means this pattern may include various possible combination of component excluding RO, here it can be
RABC + other, or a combination of RB, RAB, RBC.

### 2.5. Differential gene expression analysis

We used Limmar R package(Ritchie et al., 2015) to perform differential gene expression analysis between two target clusters. Then differential expressed genes were identified as those met these criteria: 1) FDR adjusted p value of F test < 0.01; 2) the absolute value of log2(fold change) were larger than 2.

### 2.6. Hierarchical clustering of the number and expression of DEGs among 9 patterns

We figured out the specific number of differential genes between two clusters by differential expression analysis between each pair patterns. Then the resulting count matrix was clustered using Ward.D2 by pheatmap R packages(Kolde, 2012). Similar method was applied to research *CD45* isoform pattern with the DEGs expression.

### 2.7. Developmental trajectory inference

We applied Monocle (version 2)(Trapnell et al., 2014) algorithm with the signature genes of different functional clusters and junctions representing CD45RO and CD45RA to order CD8+ T cells excluding MAIT cluster in pseudo time. We first converted TPM value into normalized mRNA counts by the “relative2abs” function in monocle and created an object with parameter “expressionFamily = negbinomial.size” according to the Monocle tutorial. Then the trajectory was determined by the default parameters of Monocle.

### 2.8. TCR sharing analysis

The TCR sequences of each single T cell were provided by Zheng et.al(Zheng et al., 2017). Each unique alpha-beta pair was defined as a clonotype. If one clonotype was present in at least two cells, cells harboring this clonotype would be considered as clonal. Then, the number of cells with such alpha-beta pair indicated the degree of clonality of the clonotype. The number of shared clonotypes across five types of T cell, including Tnaive, Tcm, Tem, Temra, Tem.ex (Tem in exhausted clusters) was calculated then plotted.

## 3. Results

### 3.1. The distribution of CD45 isoforms of T cell in tumor microenvironment

In order to identify the isoform composition of *CD45* across T cells in different state accurately, we utilized the expression of seven junctions to represent CD45 isoforms (Fig. 1A). CD45RA+ mainly appeared in naïve and effector T cells, while not in exhausted T cells and Tregs (Fig. 1B and C). However, the expression of CD45RO isoforms showed the opposite pattern with CD45RA+ isoform (Fig. 1D). Meanwhile, Tregs showed the highest level of CD45RO expression followed by Thelp and cytotoxic T cells. The distribution of CD45RO isoform in peripheral blood appeared in a lower proportion than normal and tumor tissue across the five patients (Fig. 1E and F). Thus, the distribution of CD45 isoforms depend on tissue resource, cell type, and functional state.

### 3.2. The functional difference of T cell with various CD45 isoforms

To clarify the influence of CD45 isoforms on T cell phenotype, we applied a new pattern analysis to seven junctions of CD45. Firstly, we performed pattern analysis to assign all cells into 9 new clusters with the biased expression of seven junctions (Fig. 2A). Surprisingly, eleven clusters identified by all the gene expression profile by Zheng et al(Zheng et al., 2017), possessed multiple patterns, indicating the high heterogeneity within clusters, regardless of tissue resource (Fig. 2B). We hierarchical clustered these 9 patterns both by the differential gene number and the expression of differential genes. Then,we found that T cell may be grouped into 3 populations: CD45RO^high^ T cells (pattern 5,6), CD45A^high^ (from CD45RABC) T cells (pattern 2,3,1), and CD45RB^high^ CD45RO^low^ T cells (pattern4,9,7,8)across all T cells (Fig. 2C and D). CD45RO^high^ T cells highly expressed exhaustion markers *PDCD1*, *CTLA4* and lowly expressed naïve and effector markers *GNLY*, *LEF1*, *CCR7*, opposite with CD45RA^high^, while CD45RB^high^ CD45RO^low^ T cells showed the intermediate state (Fig. 2D). In brief, although there are multiple *CD45* isoform patterns in T cells, CD45RO and CD45RA plays the dominated role in T cell phenotype.

**Figure 2.**
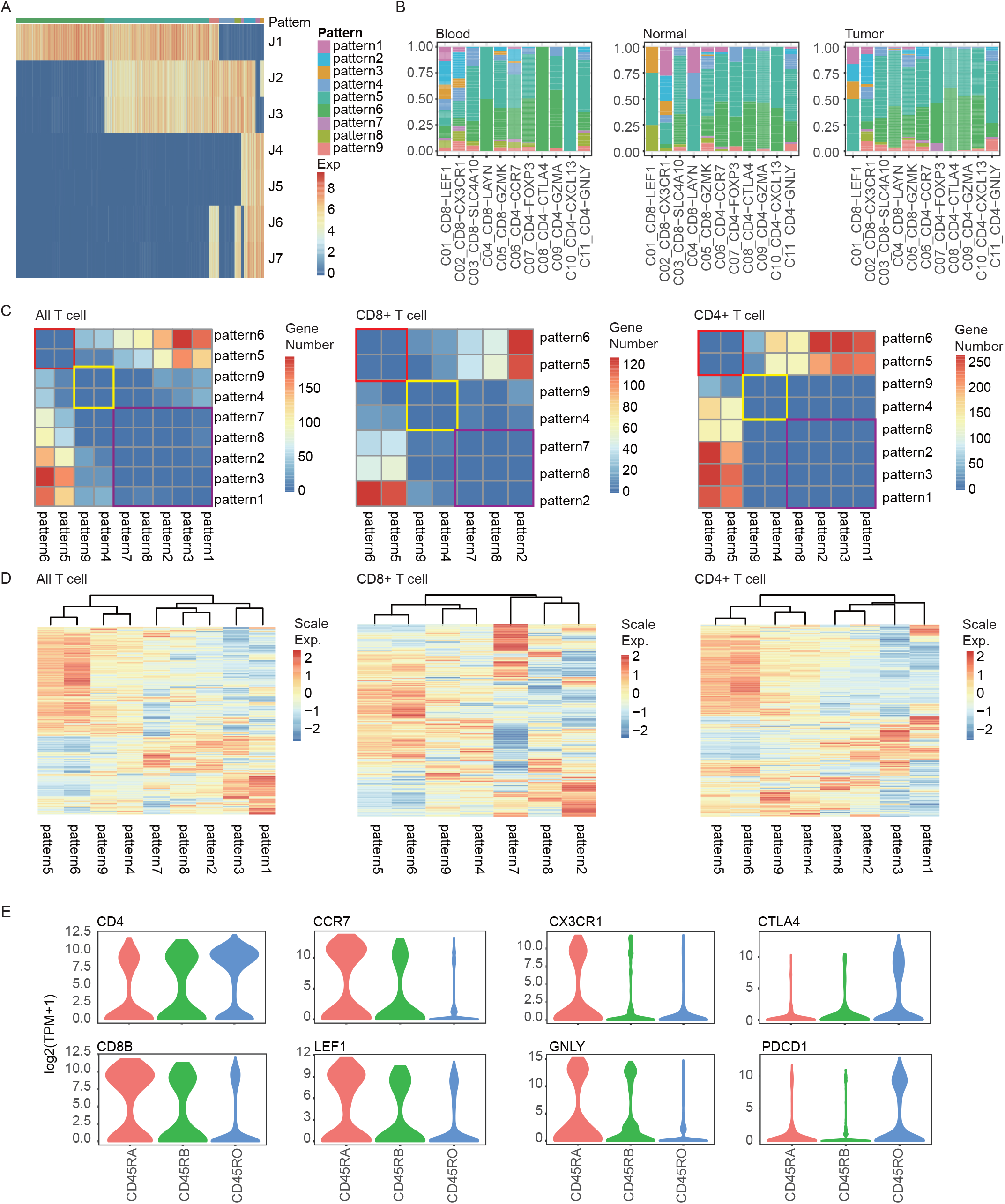
The functional difference of T cells with various CD45 isoforms. (A) The expression heatmap of junctions related with five CD45 isoforms (see FigureS5 A). Information of patterns with different CD45 isoforms composition is colored for each cell. (B) The fractions of nine patterns defined in T cells in each cluster across different tissue resources, including blood, normal tissue and tumor tissue. C, Upper, The heat map of DEGs number between pairwise patterns across T cells, CD8+ T cells, and CD4+ T cells. Lower: z-score normalized mean expression of all DEGs in each pattern across T cells, CD8+ T cells and CD4+ T cells. (D) Violin plots show the expression difference in CD45RAhigh, CD45ROhigh and CD45RBhigh subpopulation.

### 3.3. The sub-clusters and novel developmental subsets connectivity determined by CD45 isoforms

To explore the sub-clusters will promote our understanding about the heterogeneity in T cells. We used the widely accepted method which use the expression of CD45RA and *CCR7* to separate CD8+ T cells into four types: Tnaive,Tcm,Tem, and Temra. (Fig. 3A). Cells in cluster C03, C04 and C05 clusters mainly were Tem, while cluster C01 and C02 showed relative high heterogeneity (Fig. 3B). Cells in Tnaive and Tcm showed high expression of naïve markers, such as *LEF1* and *SELL*. But Tcm showed lower expression of these naïve markers than Tnaive, indicating Tcm was in the state of transition and biased to resting state. Tem moderately expressed the effector and exhausted markers, such as *GNLY*, *CX3CR1*, *CTLA4*, and *PDCD1*, implying Tem was also in the transition state but biased to activation state. Temra were characterized by the highest expression of effector and cytotoxic genes such as *NKG7* and *GNLY*. Tem in exhausted clusters (C04_CD8.LAYN, Tem.ex) highly expressed *CTLA4* and *PDCD1* (Fig. 3C). Pseudotime analysis indicated that there were two distinct developmental trajectories. Specifically, the first relative process began with Tnaive, followed by Tcm, Tem, and ended in Tem in exhausted state. The second process began with Tnaive, followed by Tcm, Tem, and ended in Temra (Fig. 3D). These two trajectories revealed the T cell state transition from activation to exhaustion and T cell state from memory to re-activation. What’s more, the clonal analysis based on those identical TCRs from common ancestors also proved this finding (Fig. 3E). In conclusion, we determined two subpopulations in state of transition, Tcm and Tem. In addition, we uncovered a novel development trajectory of tumor infiltrating T cell from Tcm to Temra.

**Figure 3.**
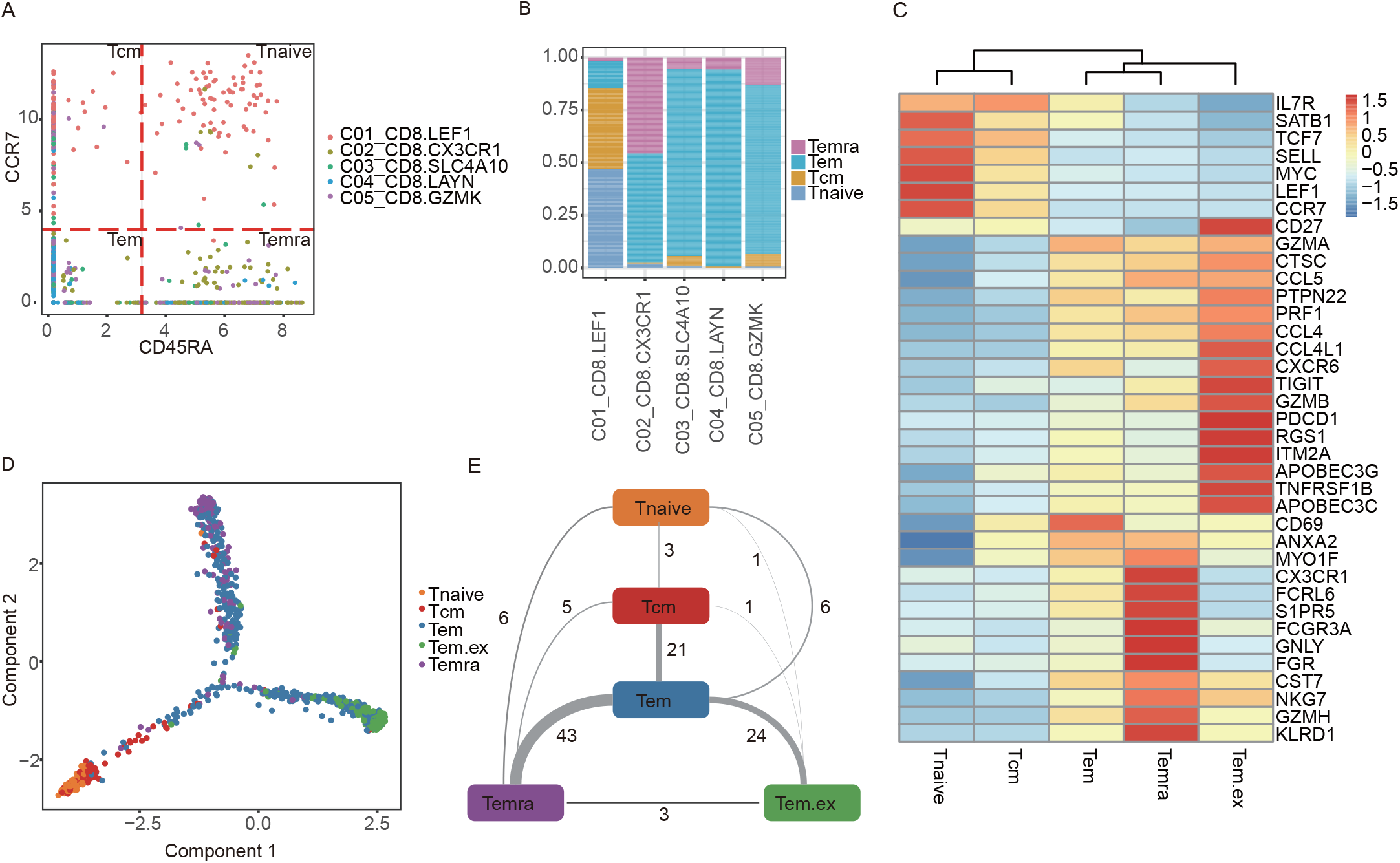
The sub-clusters and novel developmental subset connectivity determined by CD45 isoforms. (A) The expression of CD45RA and CCR7 across CD8+ T cells. Four types of T cells (Tnaive, Tem, Tcm, and Temra) are defined by the expression of CD45RA and CCR7. The thresholds of CD45RA and CCR7 were determined by the normalized count distribution. (B) The fractions of four types defined in CD8+ T cells in each cluster. (C) Z-score normalized mean expression of all DEGs in five types (Tem in C04_CD8.LAYN defined as Tem.ex). (D) CD8+ T cells (excluding MAIT cells) were ordered along pseudo time in a two-dimensional state-space defined by Monocle2. each point with different colors corresponds to individual cells in different types. (E) Cell state transition of CD8+ T cell types inferred by shared TCRs. Lines connecting different clusters are based on the degree to TCR sharing, with the thickness of lines representing the number of shared TCRs.

### 3.4. The potential clinical value of Temra

To make a clear relationship between the five types CD8+ T cells and clinical information, we analyzed their distribution features across different patients and tissue resource. Tnaive and Tcm were mainly present in peripheral blood, while Tem was mainly in tumor adjacent liver tissue. We also noticed a trend of decreased CD8+ Temra cells in HCC from late stage patients compared with early stages, while CD8+ Tem cells in exhausted state showed the opposite(Fig. 4A). At the meantime, the percentages of CD8+ Temra cells were decreased significantly in tumor tissue than in normal tissue (Fig. 4B). Thus, these results indicated that Temra in CD8 T cell were in effector and cytotoxic state and may be used for clinical diagnosis and cancer targets.

**Figure 4.**
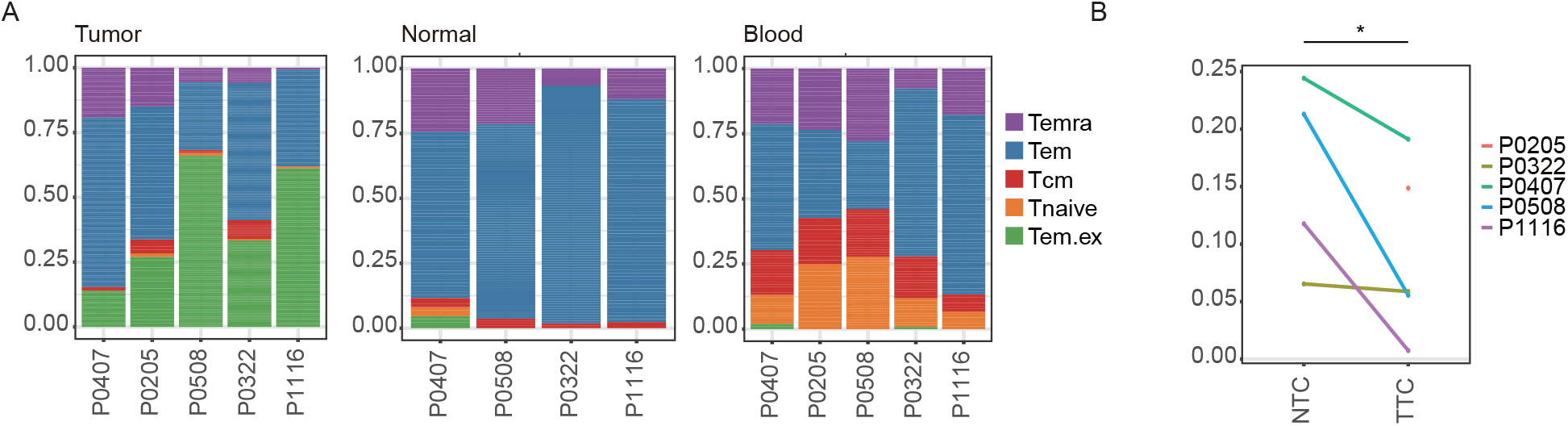
The potential clinical value of Temra. (A) The fractions of five types defined in CD8+ T cells in each patient across peripheral blood, adjacent normal, and tumor tissues. (B) The percentages of Temra in adjacent normal tissues, tumor tissues.

## 4. Discussion

Our study systematically evaluated the distribution characterization of CD45 isoforms across different functional T cells from six hepatocellular carcinoma patients at single-cell level. High level of CD45RO frequency was observed in exhausted T cells and tumor-infiltrating Treg while low in naïve and effector T cells. Increasing of exhausted T and tumor-infiltrating Treg were certainly related with worse prognosis(Jiang, Li, & Zhu, 2015; Piersma et al., 2007). However, previous studies have demonstrated that CD45RO+ memory T lymphocyte infiltration led to a favorable clinical outcome in solid tumors(Galon et al., 2006). We supposed it is the cell heterogeneity that contribute to this contradictory phenomenon. That is to say, the wide distribution of CD45RO+ cells among various T function subpopulations led to the uncertain of the dominated cell type in bulk RNA-Seq. Thus, it is essential to make it clear how T cell heterogeneity affects the prognosis at isoform levels.

The isoforms of CD45 displayed different distribution among different T cell populations. To further understand the relationship between isoform component and the state of clusters, we creatively adopt the pattern analysis of CD45 junction expression. Surprisingly, we found that various CD45 isoforms coexist in most T cell from different clusters or tissue resource, which has not been described before. After differential expressed gene analysis between different patterns, we found all the cells are separated into three main groups, CD45RO^high^ T cells, CD45RA^high^ T cells and CD45RB^high^ CD45RO^low^ T cells. Among the three groups, CD45RO^high^ population showed the opposite gene expression profiles with CD45RA^high^ population. CD45RB^high^ CD45RO^low^ population was in the middle state and owned the minimum cell compared with CD45RO^high^ and CD45RA^high^. In addition, the expression of CD45RO was negatively connected with CD45RA significantly. A plausible explanation is that the key isoforms of CD45 dominate the function and state of cells, no matter how many kinds of CD45 isoforms coexists in a cell. Specifically, it is CD45RO and CD45RA that may dominate the influence of CD45. The pattern analysis that we proposed inspires people the method development to research cell phenotype and alternative splicing at the single-cell level.

It has widely known that the expression of CD45RA+ and *CCR7* separate CD8+ T cells into four types: Tnavie, Tcm, Tem, and Temra. In our study, we succeed to obtain these four CD8+ subpopulations and expression profile of these clusters. The developmental trajectory of these clusters was discovered by the pseudo time analysis and confirmed by TCR analysis. We found a novel potential development direction from Tcm to Temra considering CD45RA expression, failing to be detected by the previous research that just applied the expression of genes to pseudo time. We also noticed two cell subsets in state of transition, Tcm and Tem out of exhausted state, that was not identified by Zheng.et al(Zheng et al., 2017). Thus, this emphasizes it is important to take advantage of the expression of isoforms to detect novel cell subsets and novel development direction.

Temra was in the terminal period of developmental trajectory and showed the highest expression of effector and cytotoxic genes, including *NKG7*, *GNLY*, and *CX3CR1*. This performance was in accordance with the previous result based scRNA-seq(Szabo et al., 2019). Meanwhile, the ratio of Temra in tumor was inversely proportional to tumor stage, implying the potential clinical application of Temra. However, this outcome needs more samples to verify due to limited samples in this study. In summary, systematical analysis of CD45 isoforms promotes our understanding about T cell heterogeneity at the level of alternative splicing. These results inspire more researches on the role of alternative splicing in T cell function heterogeneity and cancer immunotherapies.

## Conflict of interest

All authors declare no conflicts of interest

## Acknowledgement

This study is supported by Science, Technology and Innovation Commission of Shenzhen Municipality under grants (No. GJHZ20180419190827179 and No. JCYJ2017041253248372). We also thank Peking University BIOPIC Data Access Committee (PUBDAC) for approving downloading raw single-cell RNA data of HCC as well as Si Qi for her proper guide about article writing.

